# Multiple sources of uncertainty confound inference of historical human generation times

**DOI:** 10.1101/2023.02.23.529751

**Authors:** Aaron P. Ragsdale, Kevin R. Thornton

## Abstract

Wang *et al*. (2023) recently proposed an approach to infer the history of human generation intervals from changes in mutation profiles over time. As the relative proportions of different mutation types depend on the ages of parents, binning variants by the time they arose allows for the inference of average paternal and maternal generation intervals over times. Applying this approach to published allele age estimates, Wang *et al*. (2023) inferred long-lasting sex differences in average generation times and surprisingly found that ancestral generation times of West African populations remained substantially higher than those of Eurasian populations extending tens of thousands of generations into the past. Here we argue that the results and interpretations in Wang *et al*. (2023) are primarily driven by noise and biases in input data and a lack of validation using independent approaches for estimating allele ages. With the recent development of methods to reconstruct genome-wide gene genealogies, coalescence times, and allele ages, we caution that downstream analyses may be strongly influenced by uncharacterized biases in their output.

---

Recent years have seen the rapid development of methods for reconstructing genetic genealogical structures of large unrelated cohorts (Speidel *et al*., 2019; Wohns *et al*., 2022; Hubisz *et al*., 2020), which are comprised of a series of gene genealogies (or trees) along the genome. Reconstructed genealogies (or informative summaries of them) have the potential to transform population genetic inference. Biological and evolutionary processes determine the structure of these genealogies, including the distribution of mutations. In particular, the age of a variant can be estimated by mapping its mutation to the branch of the genealogy consistent with the observed data.

The past few years have also seen efforts to sequence large sets of pedigrees, providing increased resolution of important parameters of genome biology such as direct measurements of mutation rates and profiles. Through high-coverage sequencing of multiple generations of families, *de novo* mutations can be determined as maternally or paternally inherited. With a large enough sample size (meoises), both the number of mutations and proportions of mutation types (i.e., the *mutation spectrum* of A→C, A→G, etc. mutation types) can be correlated with parental age and sex (JÓnsson *et al*., 2017; Halldorsson *et al*., 2019). These two sets of inferences, the estimated ages of mutations and the parental age- and sex-dependence of the mutation spectrum, can be combined to infer the history of average maternal and paternal generation intervals for human populations of diverse ancestries (MaciÀ *et al*., 2021; Wang *et al*., 2023). In order to avoid overfitting, this approach requires making a number of assumptions about the constancy of the mutational process over time and its similarity across populations (Harris, 2015; Mathieson and Reich, 2017; Harris and Pritchard, 2017; DeWitt *et al*., 2021), and of negligible effects of selection and biased gene conversion (Lachance and Tishkoff, 2014; GlÉmin *et al*., 2015) on shaping the profile of segregating mutation types.

Gao *et al*. (2023) have recently criticized this approach. They show that observed changes in the mutation spectrum over time cannot be explained by changes in maternal and paternal generation intervals alone. Male and female generation times are required to change in different directions within the same time interval in order to explain the data for different classes of mutation. They argue that factors other than changes in generation intervals, including genetic modifiers and environmental exposure, must explain observed variation in mutation profiles. Gao *et al*. (2023) also point out that pedigree-based estimates of the *de novo* mutation spectrum do not agree with the mutation spectrum among young variants in existing population-level datasets, potentially biasing such approaches. Because such mutations occurred recently, these discrepancies would require strong mutation-type-specific selection or large recent shifts in the *de novo* mutation spectrum.

In this note, we examine both the results and conclusions of Wang *et al*. (2023). We first consider the reported inferred generation time histories and find that they are inconsistent with current understanding of human population history, in particular population structure within Africa. In exploring the source of this inconsistency, we show that allele age estimates are not just noisy, but age-stratified mutation spectra reconstructed using independent methods do not agree, with mutation profiles diverging in opposing directions. Thus, the results from Wang *et al*. (2023) do not reproduce when applying different estimates of allele ages from the same population samples. We further discuss the disagreement between the mutation rate profile found in pedigree studies and from young variants, the source of which is not well understood and complicates their comparison. In conclusion, we suggest that current methods of genealogical reconstruction and allele age estimation should continue to be evaluated for subtle biases, and that downstream analyses using estimated allele ages and mutation profiles should carefully validate their results.

## Long-lasting differences in population-specific generation intervals

Applying the proposed inference approach to multiple populations of different continental ancestries, Wang *et al*. (2023) estimated that the ancestors of European, East Asian, and South Asian populations included in the 1000 Genomes Project Consortium *et al*. (2015) dataset (1KGP) have a history of significantly reduced average generation times compared to West African populations. These differences extend to over 10,000 generations, the time period highlighted in their study. In discussing this inference the authors state, “[T]he difference among populations beyond 2000 generations ago reflects population structure in humans before their dispersal out of Africa, a structure that is not fully captured by the 1000 Genomes AFR sample. This implies that the simple labels of ‘African’ and ‘non-African’ for these populations conceal differences in generation times that existed on our ancestral continent.” Indeed, a number of recent genetic studies suggest that human population structure within Africa extending hundreds of thousands of years into the past has shaped present-day genetic variation (Hammer et al., 2011; Hsieh et al., 2016; Hey et al., 2018; Ragsdale and Gravel, 2019; Lorente-Galdos et al., 2019; Durvasula and Sankararaman, 2020).

However, in extending their analysis deeper into the past, Wang *et al*. (2023) find that ancestral generation intervals do not converge until many 10s of thousands of generations ago. Assuming an average generation time of 25–30 years, this corresponds to one to two million years in the past. This observation would require some portion of the ancestries of Eurasian and West African populations to have remained isolated for many hundreds of thousands of years, for those structured ancestral populations to have had large differences in average generation times over the course of this history, and for those groups to have contributed substantively to different contemporary human populations. While such a scenario of long-lasting isolation among ancestral populations is not impossible, it is not supported by genetic (e.g., Ragsdale et al., 2022) or archaeological (e.g., Scerri et al., 2018) evidence, which rather suggest at least periodic connectivity of ancestral human populations within Africa.

Genetic studies have estimated the Eurasian–West African divergence (i.e., the time of recent shared ancestry) at only ≈ 75ka (thousand years ago) (e.g., Pagani et al., 2015; BergstrÖm et al., 2020). While population genetic studies vary considerably in estimated split times, even those that infer deeper human divergences place the Eurasian–West African divergence at 100–150ka (e.g., Schlebusch et al., 2017). If such estimated divergence times represent the majority of ancestry of the two groups (while allowing for a smaller portion to be due to long-lasting structure), then the shared portion of ancestry should be subject to the same generation intervals prior to the divergence time. Any differences in the mutation spectrum from those epochs would be driven by differences in generation times affecting the minority of ancestry that remained isolated.

As a simple test of such a scenario, we considered a model of archaic admixture within Africa, allowing for some proportion of admixture from a diverged lineage into West African populations (such as ≈ 10% as inferred by Durvasula and Sankararaman (2020), Figure 1). At 10,000 generations ago, average paternal and maternal generation intervals in the ancestors of Eurasians were inferred to both be ≈ 20 years, while the ancestral African generation intervals were at least 28 and 23 (see Figure S4 in Wang *et al*. (2023)). Using the same mutation model (JÓnsson et al., 2017), we can determine the generation intervals in the isolated lineage that is needed to result in a mutation spectrum matching that of the inferred generation times (see Supporting Information).

**Figure 1:**
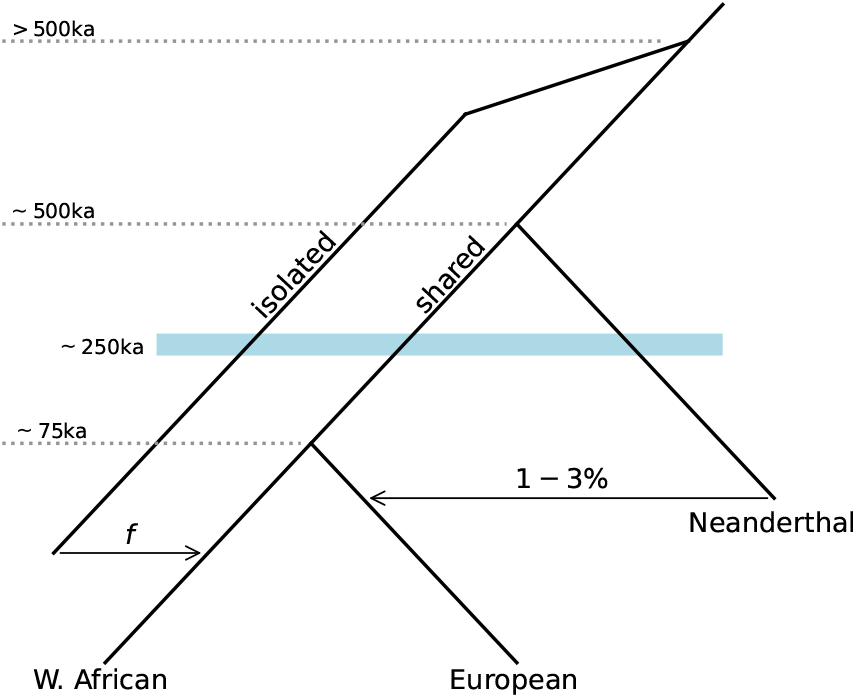
A model for archaic admixture within Africa. In such a model for “ghost” archaic introgression within Africa (Durvasula and Sankararaman, 2020), an isolated lineage diverged prior to the human-Neanderthal split and more recently contributed ancestry (with proportion *f*) to West African, but not European populations. The mutation spectrum of West African populations among alleles dating to ~ 250 – 300ka is due to a combination of mutations from the “shared” and “isolated” lineages, while the European populations’ spectrum from this time is due to a combination from the “shared” and Neanderthal branches. Wang *et al*. (2023) showed that masking Neanderthal segments in Eurasian populations had a negligible effect on their inferred generation time histories, so we ignore this small contribution when comparing observed mutation spectra between West African and Eurasian groups.

We assume that admixture proportions from the diverged and Eurasian-shared lineages (*f* and 1 – *f*, respectively) result in surviving variation from those epochs having similar proportions of contributions (in effect, ignoring differences in total mutation rate and demographic effects that may distort the magnitude of the contributed mutation spectra, see Supporting Information). With 10% admixture into the ancestors of West Africans from a diverged lineage, the mean paternal age of conception would need to be 92 and the mean maternal age 48. These are unreasonably long generation times for *Homo* species. With 20% admixture from this diverged lineage (which is larger than has been proposed or inferred in previous genetic studies), mean ages would still need to be 58 and 34. For the paternal average generation time to be less than 40, this model would require *f* = 0.4 or that the inferred generation times in the shared branch were considerably higher (Table S1).

Therefore, even assuming a model of long-lasting population structure with strict isolation within Africa, we find the reconstructed generation time intervals over the past 10,000 years from Wang *et al*. (2023) to be incompatible with plausible life histories of early humans. Given this, it is natural to ask what may be causing such mis-inference. Below we show that multiple sources of uncertainty, namely noise and bias in allele age inference and inconsistencies in trio-base estimates of mutation profiles, confound inferences of generation times from age-stratified mutation spectra.

## Inconsistencies in inferred mutation spectra over time

Wang *et al*. (2023) used published allele ages from `GEVA` (Albers and McVean, 2020) to construct mutation spectra within time periods. `GEVA` estimates allele ages by considering the number of mutations that have accumulated on the ancestral haplotype carrying the focal variant, as well as the effect of recombination in reducing the size of that ancestral haplotype. Singletons are excluded from analysis and are not assigned an age. Partitioning variants by their estimated ages shows that the mutation spectrum has changed over time, assuming that the observed spectrum of segregating variation is not biased with respect to the spectrum of *de novo* mutations occurring during that time (Figure 2A and see Figure 1C in Wang *et al*. (2023)).

**Figure 2:**
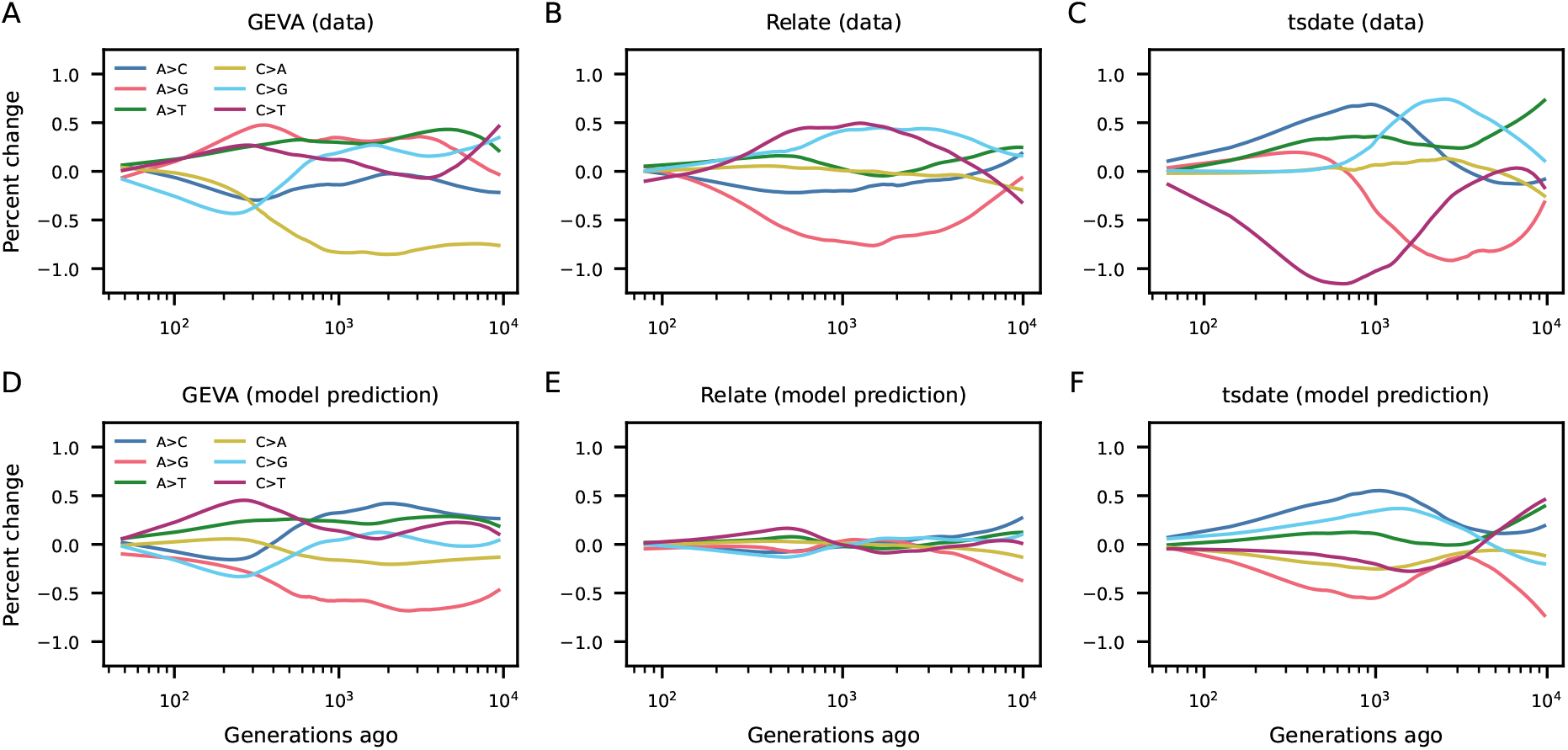
Time-stratified mutation spectra are inconsistent and poorly fit by historical generation time changes. (A-C) Three methods for estimating allele ages provide incongruous mutation spectrum histories. (D-F) In fitting historical generation intervals to each observed mutation spectrum profile, none of the mutation spectrum histories observed in the data are recovered by the predicted spectra using the inferred generation time histories. Figures S11–S13 show the mutation spectrum histories from data (top row) overlayed with predictions (bottom row). Trajectories are smoothed using LOESS regression.

In fitting generation time histories to data from the past 10,000 generations, we find that the inferred generation times provide a poor fit to the data (Figure 2D). The relative proportions of the predicted mutation spectrum trend in opposite directions for some mutation classes, confirming the concern from Gao *et al*. (2023) that a single generation time history cannot simultaneously explain each of the six mutation class frequency changes. For alleles older than 10,000 generations, the mutation spectrum fluctuates by large amounts (Figure S4). While estimated allele ages from the distant past have decreased accuracy (Albers and McVean, 2020; Wang *et al*., 2023), the large differences in proportions suggests a bias in `GEVA`’s reported dates that correlates with mutation type.

Accurate dating of variant ages is central to the inference of generation intervals from time-stratified mutation spectra. Given the poor fit of the model to the `GEVA` data and the known uncertainty in age estimation for older variants, we attempted to reproduce the inferred generation time histories using allele age estimates from independent sources, `Relate` (Speidel et al., 2019) and `tsdate` (Wohns et al., 2022), both state-of-the-art genealogical reconstruction methods able to scale to modern sample sizes. The ages of variants dated by at least two of the methods are only moderately correlated (`GEVA` and `Relate`: *r*^2^ ≈ 0.28, `GEVA` and `tsdate`: *r*^2^ ≈ 0.33, `tsdate` and `Relate`: *r*^2^ ≈ 0.64; Figure 3A-C and see Figure S20 from Wohns et al. (2022)). Imperfect correlations should be expected, as genealogical reconstruction methods have been shown to vary in the accuracy of estimated *T_M RCA_* (Brandt *et al*., 2022).

**Figure 3:**
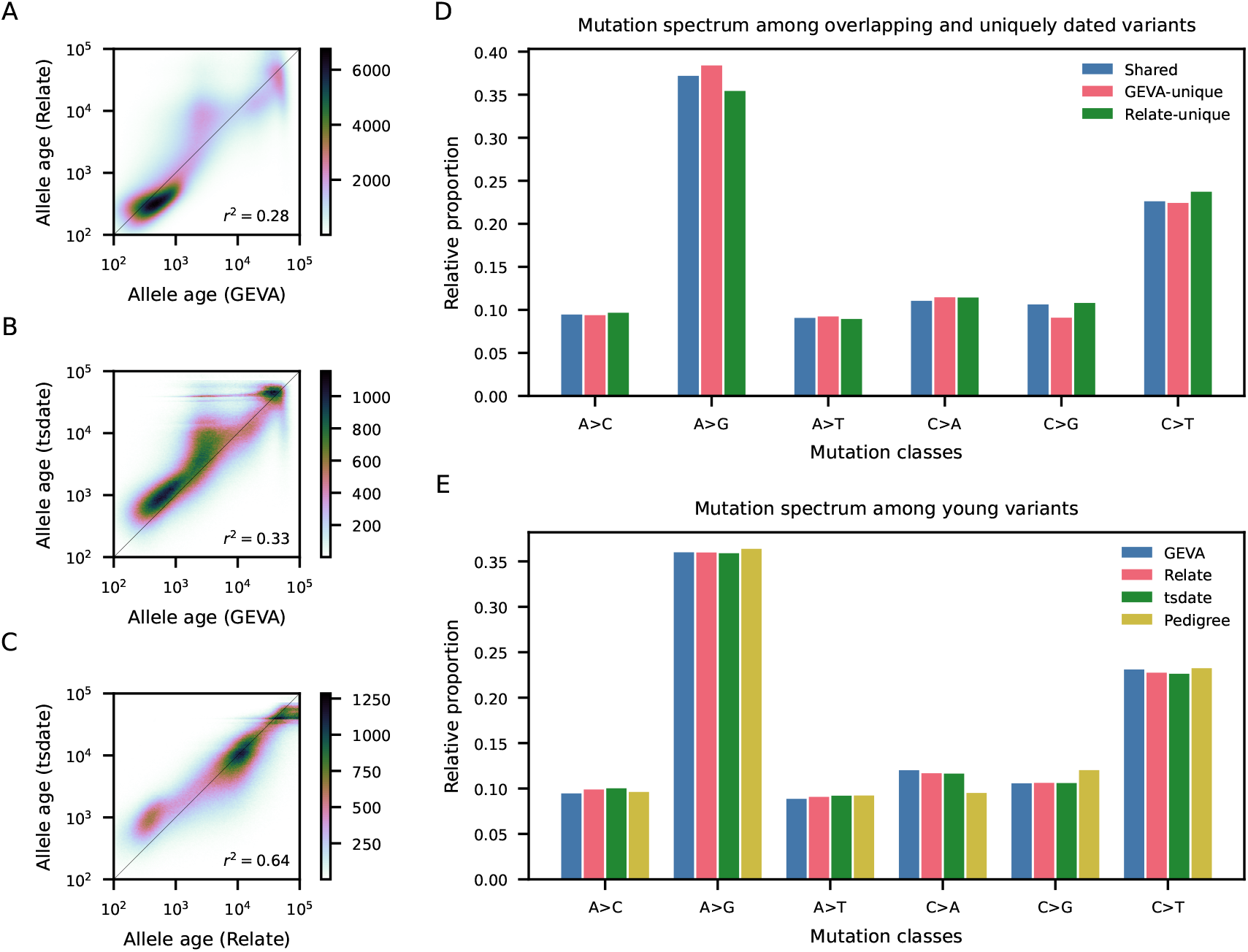
Comparing allele ages and mutation spectra across data sources. (A-C) Allele age estimates are only moderately correlated between methods. Shown here are allele age estimates for variants that were assigned ages by two or more of `GEVA, Relate` and `tsdate`. (D) The proportions of mutation types uniquely dated by `GEVA` and `Relate` differ from the variant proportions among mutations dated by both methods, indicating biases in the mutation types that are kept and discarded. Here, we compared `GEVA` and `Relate` (instead of `tsdate`) because allele ages were obtained from the same input data, namely phase three of the 1KGP. (E) While the mutation spectrum among young variants dated to < 100 generations is similar between age estimation methods, the pedigree-based estimate of the de novo mutation spectrum (JÓnsson et al., 2017) differs, in particular for C→A and C→G mutations.

Despite the imperfect correlation, it is still possible that differences in estimate allele ages are unbiased with respect to mutation type. However, we find that ages provided by each method disagree at the level of individual mutation spectrum histories (Figures 2A-C and S1–S3), with changes in mutation proportions often trending in opposite directions over the same time periods. In turn, these divergent spectrum histories provide qualitatively different inferred generation time histories (Figures S8-S10). None of them provide a reasonable fit to the data (Figure 2D-F).

Published allele ages from both Albers and McVean (2020) (`GEVA`) and Speidel *et al*. (2019) (`Relate`) used the same input dataset: the phase 3 release of 1KGP. However, they differ in the total number of variants that were assigned ages (roughly 30 million and 48 million, respectively). `GEVA` and `Relate` each discard variants that violate or are inconsistent with certain assumptions, such as allowing only a single mutation at a given site, while `tsdate` retains all data by allowing for multiple mutations at a site. Of those variants, 25 million variants were dated by both `GEVA` and `Relate`, with the remaining uniquely assigned an age by one or the other method. When comparing mutation spectrum histories restricted to variants dated by both `GEVA` and `Relate`, the two methods still predict different mutation profile trajectories (Figure S7). In comparing the proportions of mutation types assigned ages by both methods or uniquely by one, we find that those proportions differ considerably (Figure 3D). For example, `GEVA` provides age estimates for relatively more A→G mutations and fewer C→G mutations, while `Relate` provides age estimates for relatively more C→T and fewer A→G mutations. This owes to methodological differences in evaluating whether a particular variant can be reliably dated, which may have both bioinformatic (e.g., genotyping, phasing and polarization errors) and biological (e.g., recurrent mutations) causes. Therefore, when comparing mutation spectra that are conditioned on variants being dated by a particular method, it is not apparent which method, if any, provides unbiased estimates of age-stratified mutation spectra.

## Mutation spectra differ between *de novo* mutations and young alleles

The large disagreements in mutation spectrum histories between multiple variant age-estimation methods are concerning for downstream inferences that rely on them. Independent of biases in allele age estimate, there are further problems in comparing age-stratified mutation spectra to those estimated from pedigree studies (JÓnsson et al., 2017; Halldorsson et al., 2019). As Wang *et al*. (2023) acknowledge, the spectrum of *de novo* mutations identified in Icelandic trios (JÓnsson et al., 2017) differs from the spectrum of young segregating variation (e.g., variants estimated to be less than 100 generations old, Figure 3E and Table S2). Gao *et al*. (2023) argue that these differences are unlikely to be driven by biological processes.

For some mutation classes, the relative proportion of de *novo* mutations in the trio-based study differs from the young-variant spectrum by up to 0.02, which would imply a large over- or under-count of different mutation types. This is the equivalent of 10s of thousands of SNPs in the most recent bin, with the exact number depending on the variant-dating method. `GEVA, tsdate` and `Relate`, while their estimated mutation spectra differ at older times, closely agree for mutations inferred to be less than 100 generations old (Table S2). In discussing this discrepancy, Wang *et al*. (2023) state, “We found that the mutation spectrum from the large pedigree study consistently differed from the variant spectrum inferred from the 1000 Genomes Project data, possibly because we removed singletons from the polymorphism dataset to reduce errors.” However, `GEVA` does not provide estimates of allele ages for singletons, so this suggested source of discrepancy cannot be checked with their published allele ages. Both `tsdate` and `Relate` report allele ages for singletons, and their inclusion does not strongly affect the mutation spectrum for very young variants (Table S2), though it does impact the mutation profiles in older time periods (Figures S5, S6). Reported ages from `GEVA` and `Relate` both used the low-coverage phase 3 1KGP data while `tsdate` used the more recent independently sequenced high-coverage 1KGP data (Byrska-Bishop et al., 2022), so the accuracy of the mutation spectrum among young variants is unlikely to be driven by differences in coverage.

To account for the differences between the de novo spectrum from pedigree studies (JÓnsson et al., 2017) and the spectrum among young variants, Wang *et al*. (2023) subtracted this difference from each historical mutation spectrum. “[This choice] has the effect of assuming that parental ages in the pedigreed mutation dataset reflect generation times in the most recent historical bin” (Wang et al., 2023). However, the average generation time among present-day Icelanders may not reflect average generation times in worldwide populations over the past 3-5 thousand years, represented by the most recent time bin in their analysis. Still more concerning is that we do not know the source of the discrepancy between the Iceland and 1KGP mutation spectra. Without knowing this, it is unclear that simply subtracting the differences between them properly accounts for it. Instead, mutation class-specific genotyping differences may distort the underlying mutation model, potentially driving deviations in predicted mutation profiles in unexpected ways.

What could be driving the large disagreement between the spectrum of *de novo* mutations from Icelandic pedigrees and that of young variants in the 1KGP dataset? First, there could be true differences in the mutation spectrum between the Iceland cohort and 1KGP populations, although this seems unlikely, as populations of different ancestries in 1KGP have more similar recent mutation spectra to each other than to the Iceland *de novo* spectrum (including the EUR populations, Table S3). If the differences are real, then the Icelandic pedigree data is an inappropriate calibration for the mutation spectrum in other populations. Second, there could be differences in selective pressures between mutations of different classes, although selection would need to be very strong for some classes compared to others in order to see the observed difference among young variants. Third, the signal may be driven by genotyping error or bioinformatics choices, though the agreement between high- and low-coverage 1KGP datasets suggests that genotyping error does not have a strong effect in the 1KGP data. Instead, filtering and bioinformatics choices in the pedigree approach are the likely culprit (Bergeron *et al*., 2022). Indeed, Gao *et al*. (2023) show that two studies of Icelandic trios that partially overlap in families that were included (JÓnsson *et al*., 2017; Halldorsson *et al*., 2019) estimate significantly different *de novo* mutation spectra due to filtering choices (see Figure 3, supplement 2 in Gao *et al*. (2023)). Until the source of these differences are understood and properly accounted for, we caution that population-genetic inferences should avoid calibration using mutation rates and profiles from pedigree studies.

## Conclusions

Large-scale reconstruction of gene genealogies, dating mutations, and T_MRCA_ inference are all exciting developments, but these methods are quite new and will likely advance considerably in the coming years. Currently, existing methods show different accuracies even under overly-idealized conditions (Brandt *et al*., 2022). We should therefore be cautious of unexpected and subtle biases that can impact downstream analyses. In the case of generation time inference (Wang *et al*., 2023), we showed that patterns of variation that were attributed to biological processes (variation in generation times) are predominantly driven by nonbiological artifacts in the input data. As the field continues to evaluate the accuracy of allele age estimation and genealogical reconstruction methods, we recommend that analyses that rely on their output thoroughly validate their results using independent approaches.

## Acknowledgements

We thank Ziyue Gao and Priya Moorjani for helpful discussions.

