## Supporting Information for "Multiple sources of uncertainty confound inference of historical human generation times"

February 23, 2023

#### Supplemental methods

##### Generation times needed to explain long-lasting differences between populations

Using the reported generation intervals from Wang *et al.* (2023) (as shown in Figure S4 in their supplemental material), we explored scenarios that could lead to the observed differences between African and non-African populations five to ten thousand generations ago, corresponding to 150-300ka. As discussed in the main text, a 5-10 year difference in generation intervals would require long-lasting structure. Admixture from an unidentified, diverged human lineage has been proposed to explain observed genetic variation in African populations (e.g., Hey *et al.*, 2018; Durvasula and Sankararaman, 2020; Lorente-Galdos *et al.*, 2019, but see Ragsdale *et al.* (2022) for alternative models that allow for ongoing gene flow between lineages). In such models, a population that was isolated for hundreds of thousands of years contributed 5–10% ancestry to West African populations (Figure 1). The remaining 90–95% ancestry is shared between present-day Eurasian and West African populations, and this ancestry would have shared historical generation times. While other models of deep population structure in Africa have been proposed, a history of strict isolation (instead of ongoing gene flow) between diverged lineages before admixture is more likely to result in a signal of differing ancestral generation times, because ancestries and their associated generation intervals would remain distinct.

In such a scenario, differences in inferred average historical generation times between West African and Eurasian populations must be due to differences in generation times between the two diverged lineages. This is because ancestry that is shared within the common branch will have been merged and average generation intervals would have likewise been shared. Using the mutation model Wang *et al.* (2023) inferred from Icelandic pedigree data (Jónsson *et al.*, 2017), we modeled the Eurasian mutation spectrum from this time period using paternal and maternal generation times of 20 (18–22, from figure S4 in Wang *et al.*) years. The West African mutation spectrum was modeled as a mixture between this shared spectrum and the mutation spectrum from the diverged lineage, in proportions equal to the admixture proportions. This assumes (1) selection does not strongly influence mutation spectrum proportions, (2) there are no demographic effects such as severe bottlenecks that make mutation spectrum proportions unequal to admixture proportions, and (3) the rates of mutation accumulation along each lineage are similar. Additionally, age- and sex-dependent mutation rates from past populations must match the mutation model from the Icelandic trio data. It is likely that none of these assumptions perfectly hold, but these are the same assumptions in the original inference of generation time histories.

We write our mutation model as $M(p, d)$, which takes paternal and maternal ages $p$ and $d$ and outputs the expected mutation spectrum. The inferred West African-ancestral paternal and maternal generation intervals were roughly 28 and 23 years (see Figure S4 in Wang *et al.* (2023)). Then given the admixture proportion $f$ from the diverged lineage, we found generation times $p_d$ and $m_d$ in the diverged lineage such that

$$M(28, 23) = (1 - f)M(20, 20) + fM(p_d, m_d).$$

In fitting this model with $f = 0.1$ (roughly the inferred admixture proportion from Durvasula and Sankararaman (2020)), we found $p_d \approx 92$ and $m_d \approx 48$.

If we assume average ancestral paternal and maternal ages in the Eurasian-shared lineage were each 22 years, $p_d \approx 76$ and $m_d \approx 31$. With Eurasian-ancestral intervals of 22 years and $f = 0.2$ (much higher than most inferences), the paternal age would still need to be over 50 years, inconsistent with average generation times in humans and great apes. More comparisons are shown in Table S1. From this, we conclude that the generation time history inferred by Wang *et al.* (2023) is incompatible with prevailing models of deep population structure within Africa.

| Input Parameters | | | | | Fit parameters | |
|---|---|---|---|---|---|---|
| $f$ | $p_{\mathrm{EUR}}$ | $m_{\mathrm{EUR}}$ | $p_{\mathrm{AFR}}$ | $m_{\mathrm{AFR}}$ | $p_{\mathrm{d}}$ | $m_{\mathrm{d}}$ |
| 0.1 | 20 | 20 | 28 | 23 | 92.2 | 47.9 |
| 0.1 | 20 | 20 | 30 | 25 | 111.2 | 67.6 |
| 0.1 | 20 | 20 | 26 | 22 | 75.0 | 38.6 |
| 0.2 | 20 | 20 | 28 | 23 | 58.0 | 34.4 |
| 0.2 | 20 | 20 | 30 | 25 | 67.7 | 44.4 |
| 0.2 | 20 | 20 | 26 | 22 | 48.8 | 29.6 |
| 0.3 | 20 | 20 | 28 | 23 | 45.9 | 29.8 |
| 0.3 | 20 | 20 | 30 | 25 | 52.4 | 36.4 |
| 0.3 | 20 | 20 | 26 | 22 | 39.5 | 26.5 |
| 0.1 | 22 | 22 | 28 | 23 | 75.6 | 30.6 |
| 0.1 | 22 | 22 | 30 | 25 | 94.2 | 49.9 |
| 0.1 | 22 | 22 | 26 | 22 | 58.4 | 21.4 |
| 0.2 | 22 | 22 | 28 | 23 | 50.5 | 25.2 |
| 0.2 | 22 | 22 | 30 | 25 | 60.1 | 36.4 |
| 0.2 | 22 | 22 | 26 | 22 | 41.2 | 21.8 |
| 0.3 | 22 | 22 | 28 | 23 | 41.4 | 25.2 |
| 0.3 | 22 | 22 | 30 | 25 | 47.9 | 31.8 |
| 0.3 | 22 | 22 | 26 | 22 | 35.0 | 21.8 |

Table S1: **Testing admixture proportions and generation intervals under an African archaic admixture model.** In order for the generation times in the isolated branch ($p_d$ and $m_d$) to be reasonably short enough for human biology, the admixture proportion would need to be $\gtrsim 0.3$ and for the inferred generation intervals in Eurasians and West Africans to be much closer than the average values shown in Figure S4 in Wang *et al.* (2023).

#### Historical mutation spectra

We followed the filtering choices from Wang *et al.* (2023) in retaining mutations with estimated ages. Namely, triplet mutation contexts associated with a known C→T mutation pulse in Europeans (Harris, 2015) and CpG sites were removed. `GEVA` does not provide allele ages for singletons, but we considered data both with and without singletons from `tsdate`- and `Relate`-inferred ages. Variants with allele frequencies greater than 98% were removed to minimize the effect of ancestral-state misidentification.

Variants were binned by age in 100 epochs, divided such that a roughly equal number of variants fell within each bin, as in Wang *et al.* (2023). In most cases, we considered a maximum age of 10,000 generations.

Mutation profile trajectories and generation time histories were smoothed using the `loess_1d` function from the Python `loess` package, with parameters `frac=0.5` and `degree=2`.

##### Allele ages from GEVA

Allele age data from GEVA reported in ALBERS and MCVEAN (2020) were downloaded from https://human.genome.dating/download/index. We used the median joint-estimated allele ages, "AgeMedian_Jnt". To compare to allele age data estimated from Relate and tsdate, we used allele ages estimated from the Thousand Genomes Project (TGP) data source, available from http://ftp.ensembl.org/pub/grch37/release-103/variation/gvf/homo_sapiens/.

##### Allele ages from Relate

Allele age data from Relate reported in SPEIDEL *et al.* (2019) were downloaded from https://zenodo.org/record/3234689. Relate provides allele ages separately for the 26 populations in the Thousand Genomes Project. As such, we followed the approach in WOHNS *et al.* (2022) and computed the average upper and lower bounds of the edge the mutation lies over for each population it is present in. We then took the midpoint of those averaged upper and lower bounds as the allele age estimate.

##### Allele ages from tsdate

The reconstructed genealogies from WOHNS *et al.* (2022) were downloaded from https://zenodo.org/record/5512994. Data for each chromosome arm were provided in tree sequence format, and mutation ages can be estimated from the upper and lower bounds of the genealogical edge on which it arose. We kept sites with variants that were uniquely assigned to a single branch, and we considered variants that segregate among the subset of individuals from the AFR, EAS, EUR, and SAS Thousand Genomes Project superpopulations. For such sites, allele ages were found using the function `tsdate.sites_time_from_ts(ts, node_selection="arithmetic")`.

##### Reference genomes

To determine triplet contexts for each mutation, we used the reference genomes for GRCh37 (GEVA and Relate data) and GRCH38 (tsdate data). These were downloaded from http://hgdownload.cse.ucsc.edu/goldenPath/hg19/chromosomes/ and https://ftp.1000genomes.ebi.ac.uk/vol1/ftp/technical/reference/GRCh38_reference_genome/, respectively.

##### Data availability

All analyses were performed using publicly available datasets, available from the above URLs. Python scripts to run analyses described here are available at https://github.com/apragsdale/dated-mutation-spectra/

### Tables and figures

| Dataset | A→C | A→G | A→T | C→A | C→G | C→T |
|---|---|---|---|---|---|---|
| `GEVA` | 0.0946 | 0.3600 | 0.0886 | 0.1201 | 0.1057 | 0.2310 |
| `tsdate` | 0.0931 | 0.3579 | 0.0899 | 0.1146 | 0.1061 | 0.2384 |
| `tsdate` (w/singletons) | 0.0989 | 0.3598 | 0.0908 | 0.1168 | 0.1062 | 0.2275 |
| `Relate` | 0.0991 | 0.3610 | 0.0863 | 0.1124 | 0.1038 | 0.2374 |
| `Relate` (w/singletons) | 0.1002 | 0.3590 | 0.0921 | 0.1164 | 0.1060 | 0.2263 |
| Trios (phased) | 0.0953 | 0.3649 | 0.0890 | 0.0960 | 0.1216 | 0.2332 |
| Trios (all mutations) | 0.0962 | 0.3638 | 0.0923 | 0.0951 | 0.1202 | 0.2324 |

Table S2: **Mutation profiles from the past 100 generations, compared to Iceland trios.** The most recent time bin for each method included the past $\approx 150$ generations. When singletons were included (when using data from `tsdate` and `Relate`), the spectra of estimated recent standing variation were unchanged. Note that `GEVA` does not report ages for singletons. While the three methods provide similar spectra from recent mutations, the spectrum from the Iceland pedigrees differs, in particular for the C→A and C→G classes. These differences are up to 2% of the proportion among all mutations, which corresponds to an under- or over-count of up to $\sim 20\%$ of C→A and C→G mutations, respectively. This difference remains whether the spectrum is estimated from only mutations that were phased in Jónsson *et al.* (2017) or from all mutations (phased and unphased).

| Dataset | A$\to$C | A$\to$G | A$\to$T | C$\to$A | C$\to$G | C$\to$T |
|---|---|---|---|---|---|---|
| AFR (GEVA) | 0.103 | 0.354 | 0.094 | 0.127 | 0.098 | 0.224 |
| EAS | 0.111 | 0.341 | 0.103 | 0.131 | 0.094 | 0.220 |
| EUR | 0.102 | 0.355 | 0.093 | 0.125 | 0.102 | 0.222 |
| SAS | 0.095 | 0.355 | 0.090 | 0.123 | 0.099 | 0.238 |
| AFR (Relate) | 0.099 | 0.356 | 0.084 | 0.116 | 0.110 | 0.236 |
| EAS | 0.095 | 0.359 | 0.089 | 0.115 | 0.097 | 0.245 |
| EUR | 0.100 | 0.368 | 0.085 | 0.110 | 0.102 | 0.235 |
| SAS | 0.104 | 0.344 | 0.090 | 0.108 | 0.107 | 0.246 |
| AFR (tsdate) | 0.092 | 0.354 | 0.087 | 0.116 | 0.110 | 0.241 |
| EAS | 0.098 | 0.356 | 0.097 | 0.112 | 0.103 | 0.233 |
| EUR | 0.091 | 0.363 | 0.089 | 0.117 | 0.102 | 0.238 |
| SAS | 0.091 | 0.359 | 0.088 | 0.114 | 0.107 | 0.241 |

Table S3: **Mutation profiles from the past 100 generations in continental population groups.**

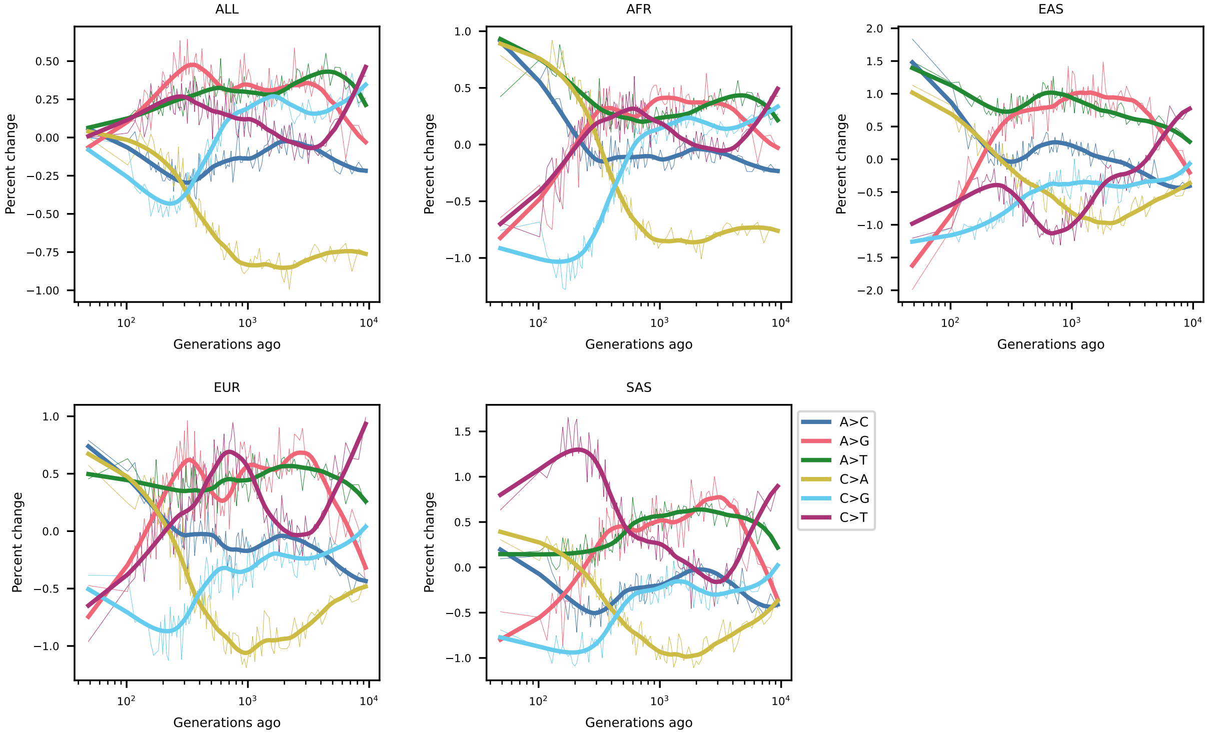


Figure S1: **GEVA-inferred mutation spectrum history.**

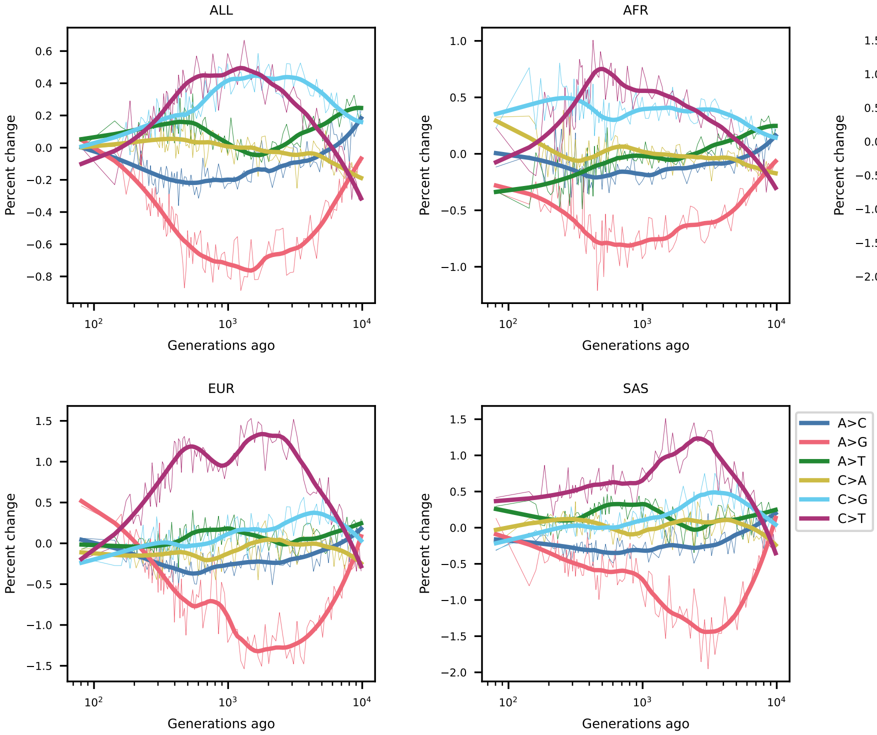


Figure S2: **`Relate`-inferred mutation spectrum history.**

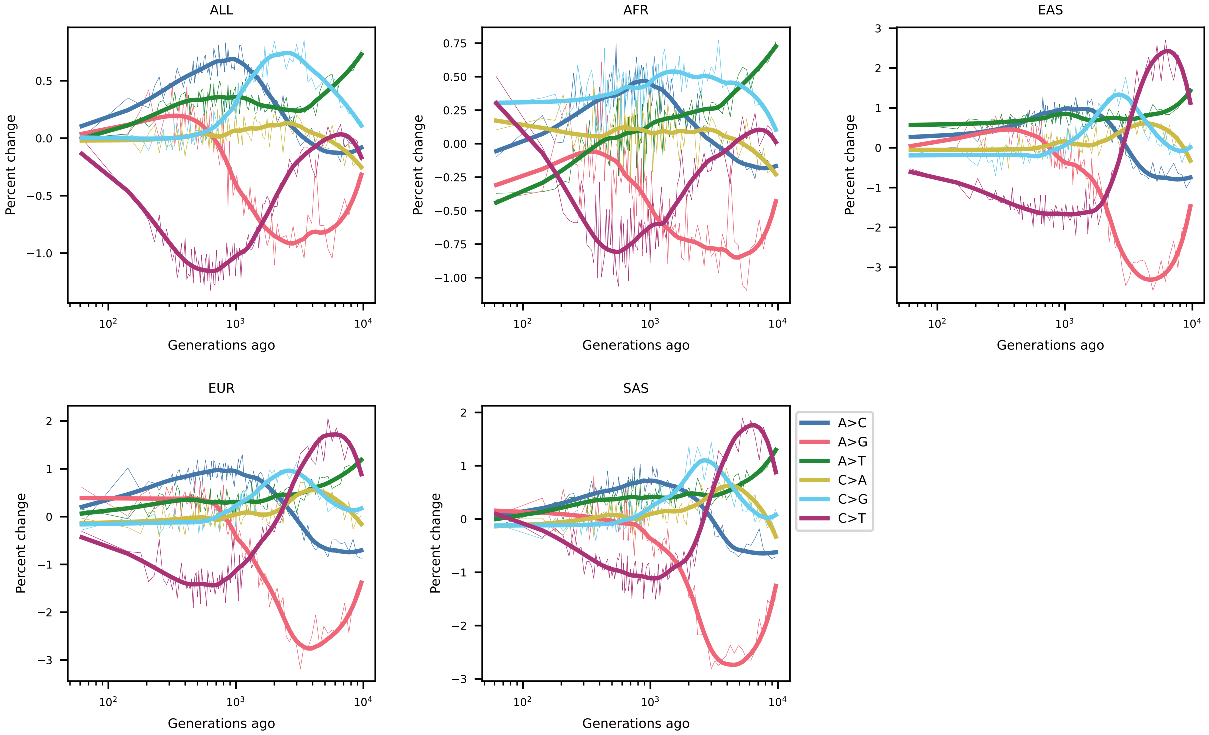


Figure S3: **`tsdate`-inferred mutation spectrum history.**

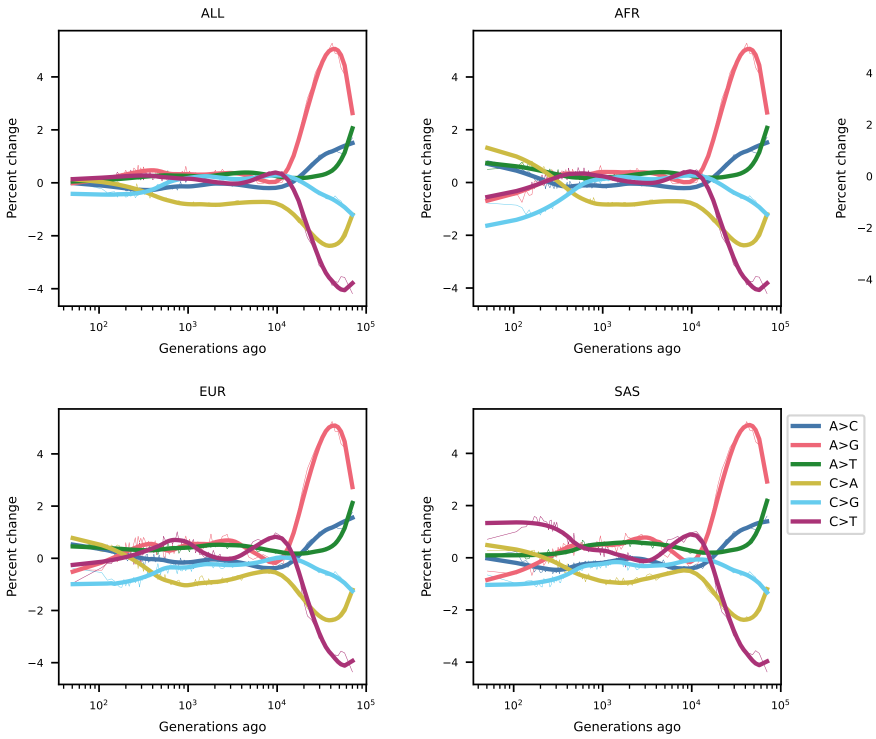


Figure S4: **`GEVA`-inferred mutation spectrum history, extending to 80,000 generations.**

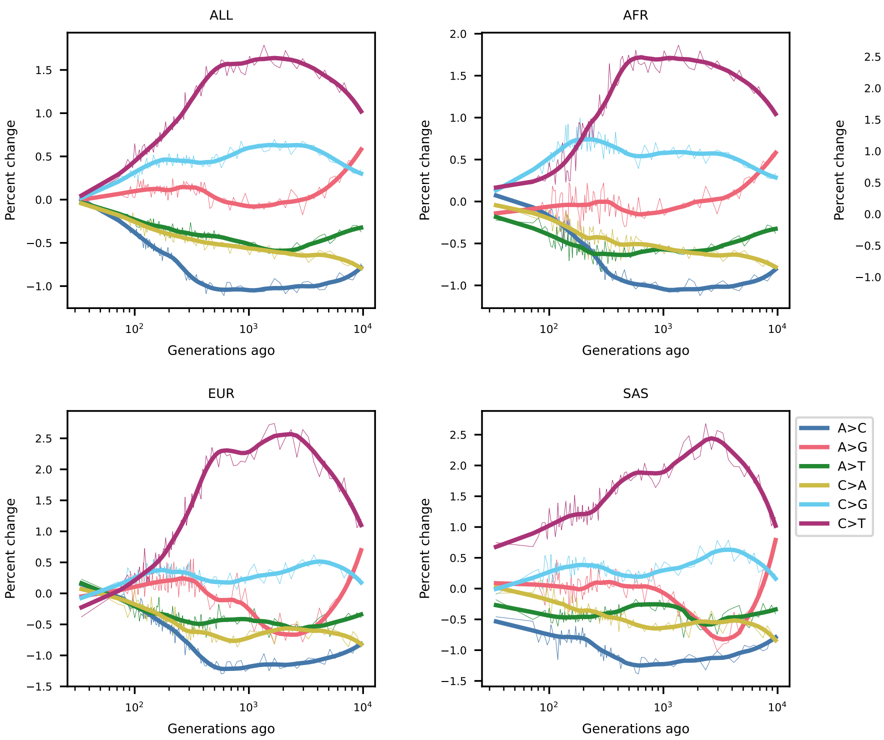


Figure S5: **`Relate`-inferred mutation spectrum history, including singletons.**

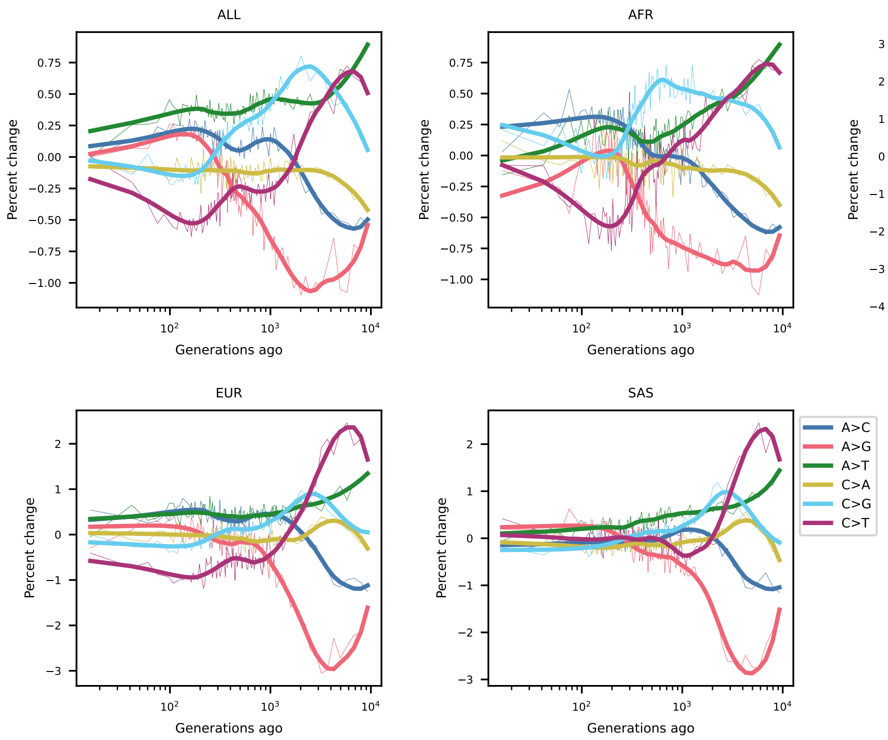


Figure S6: **`tsdate`-inferred mutation spectrum history, including singletons.**

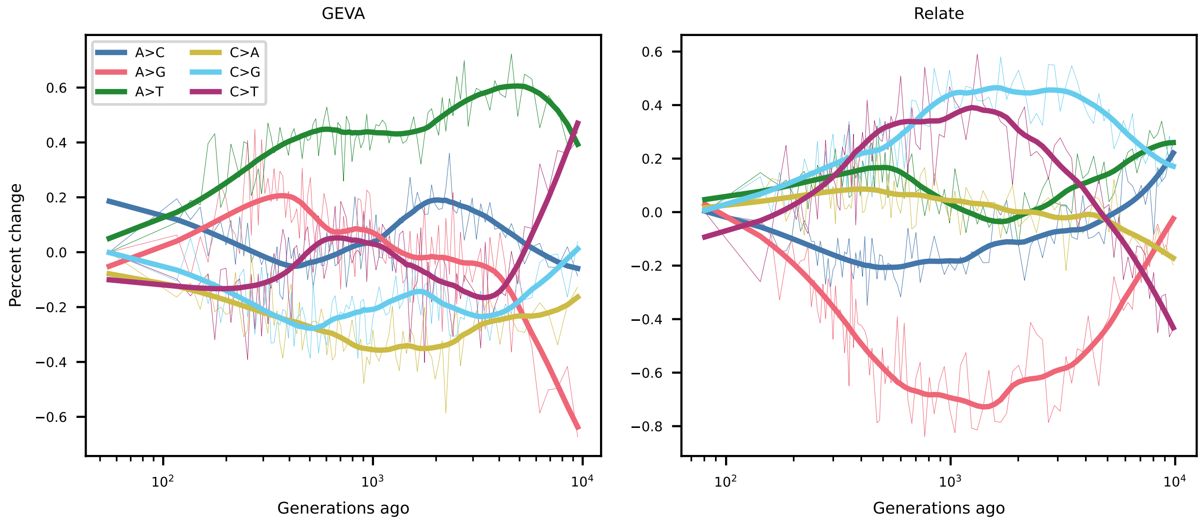


Figure S7: **Mutation spectrum histories from mutations that were dated by both `GEVA` and `Relate`.**

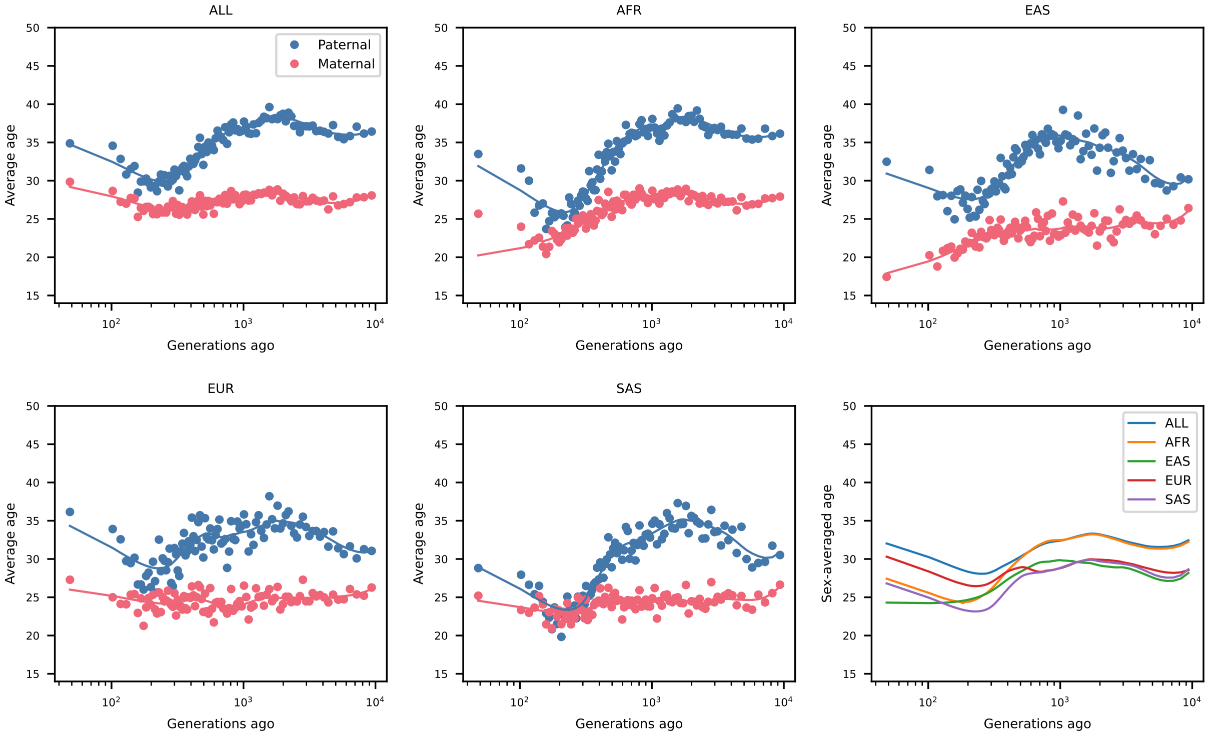


Figure S8: `GEVA`**-inferred generation time histories.**

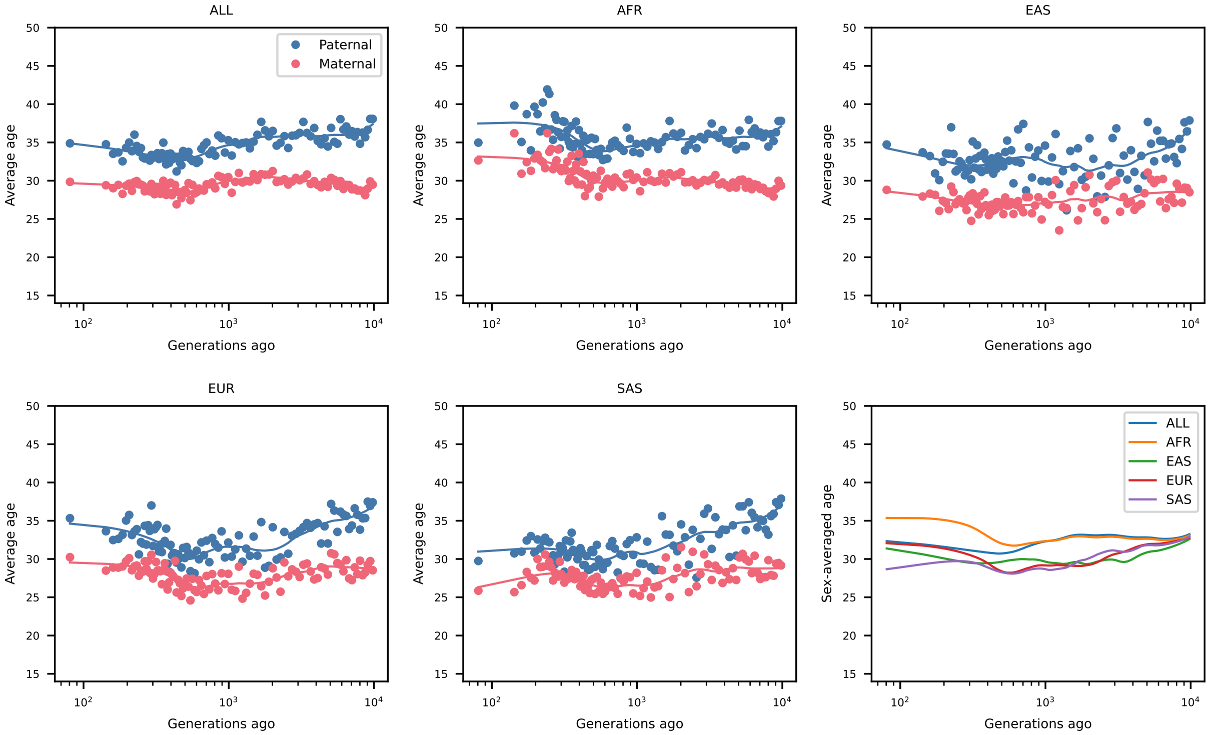


Figure S9: `Relate`**-inferred generation time histories.**

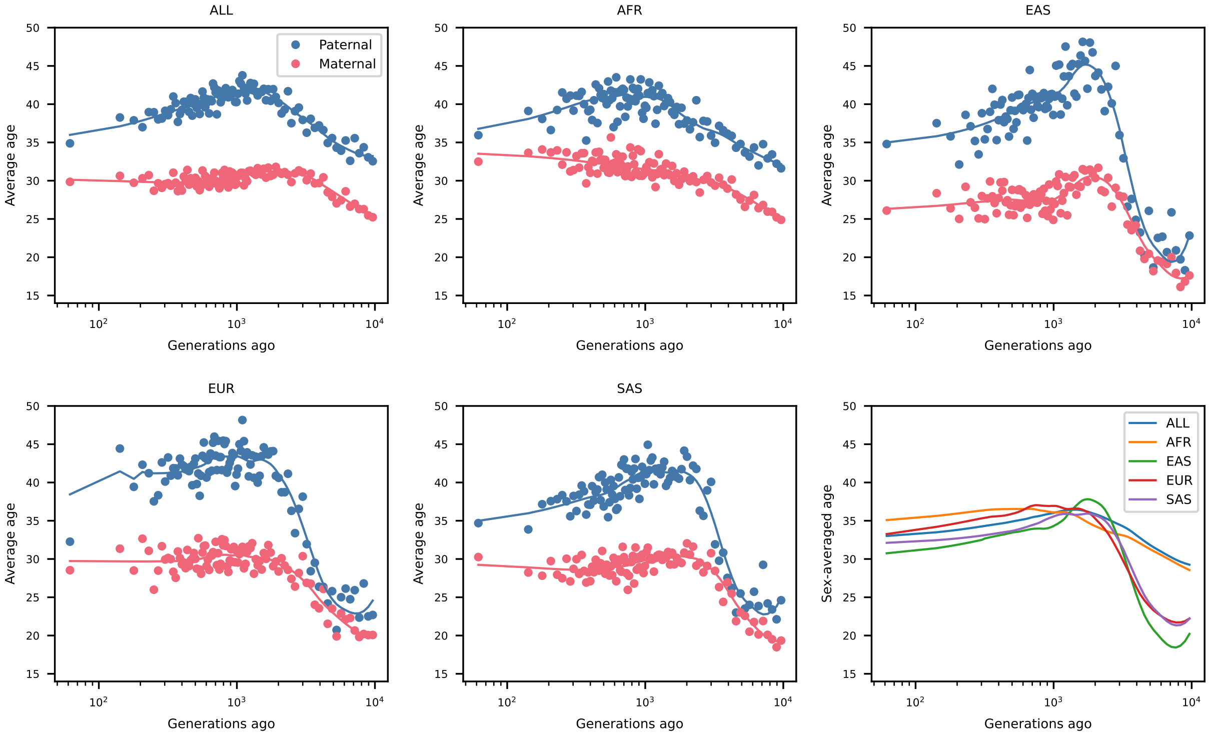


Figure S10: `tsdate`**-inferred generation time histories.**

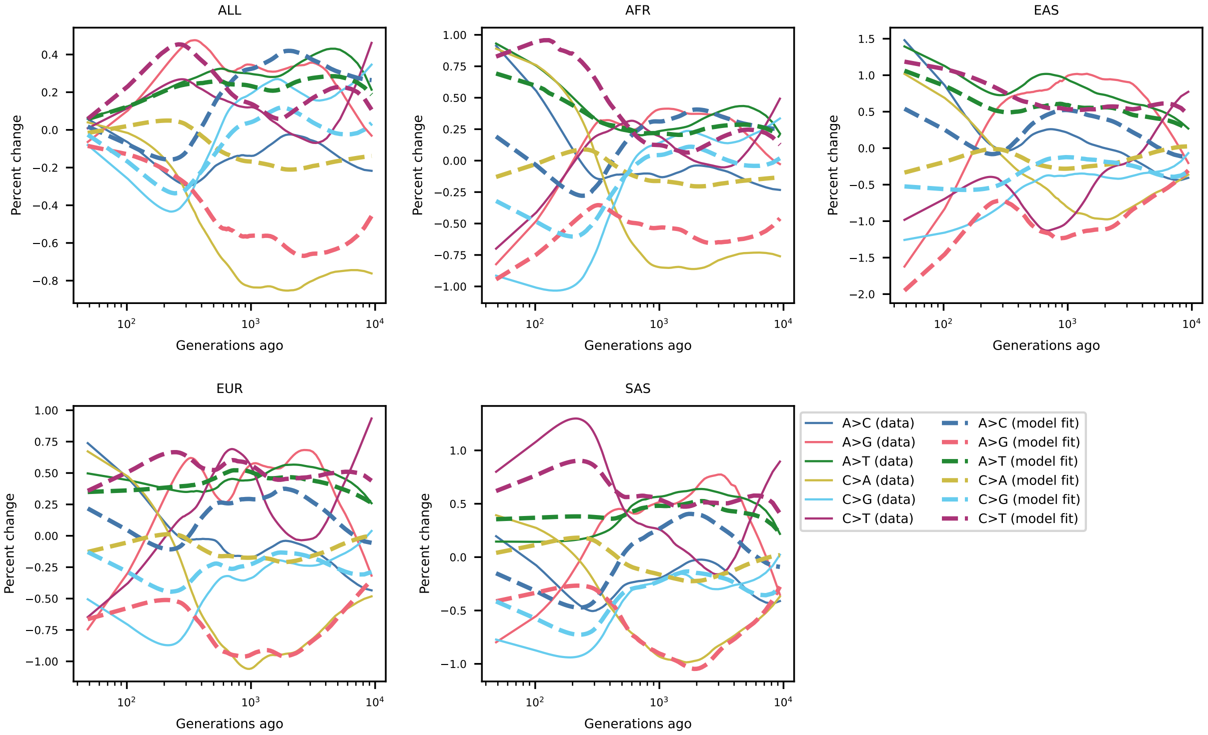


Figure S11: **Prediction of mutation spectrum history from GEVA-inferred generation times.**

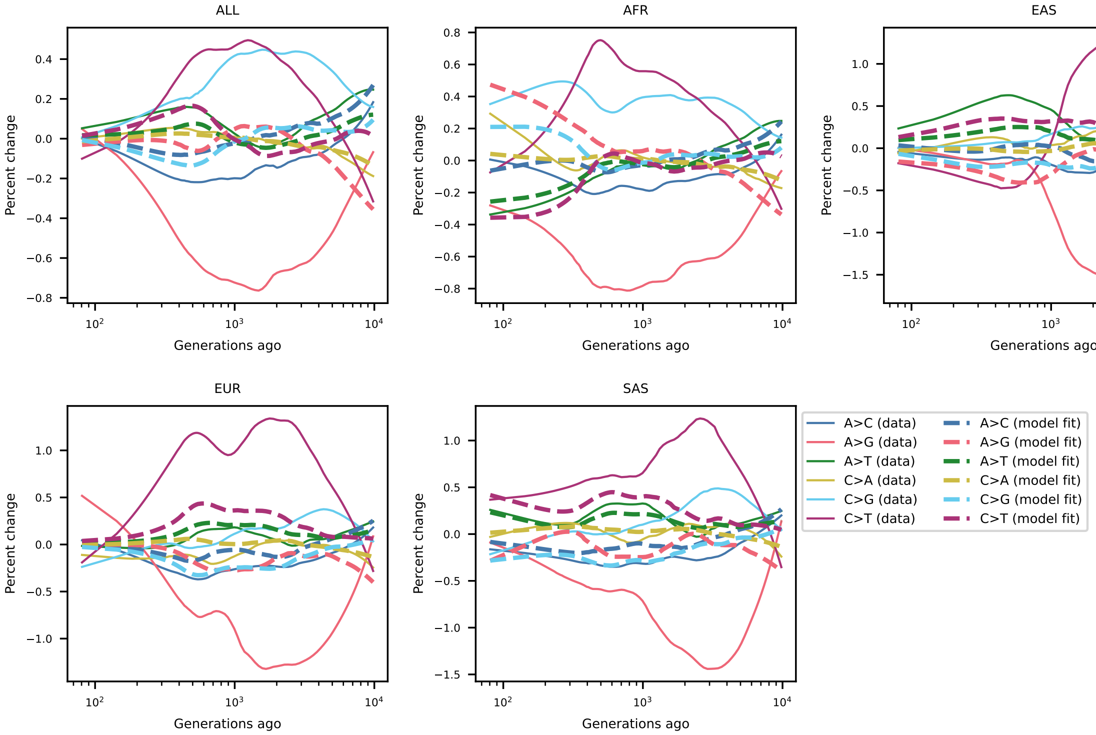


Figure S12: **Prediction of mutation spectrum history from `Relate`-inferred generation times.**

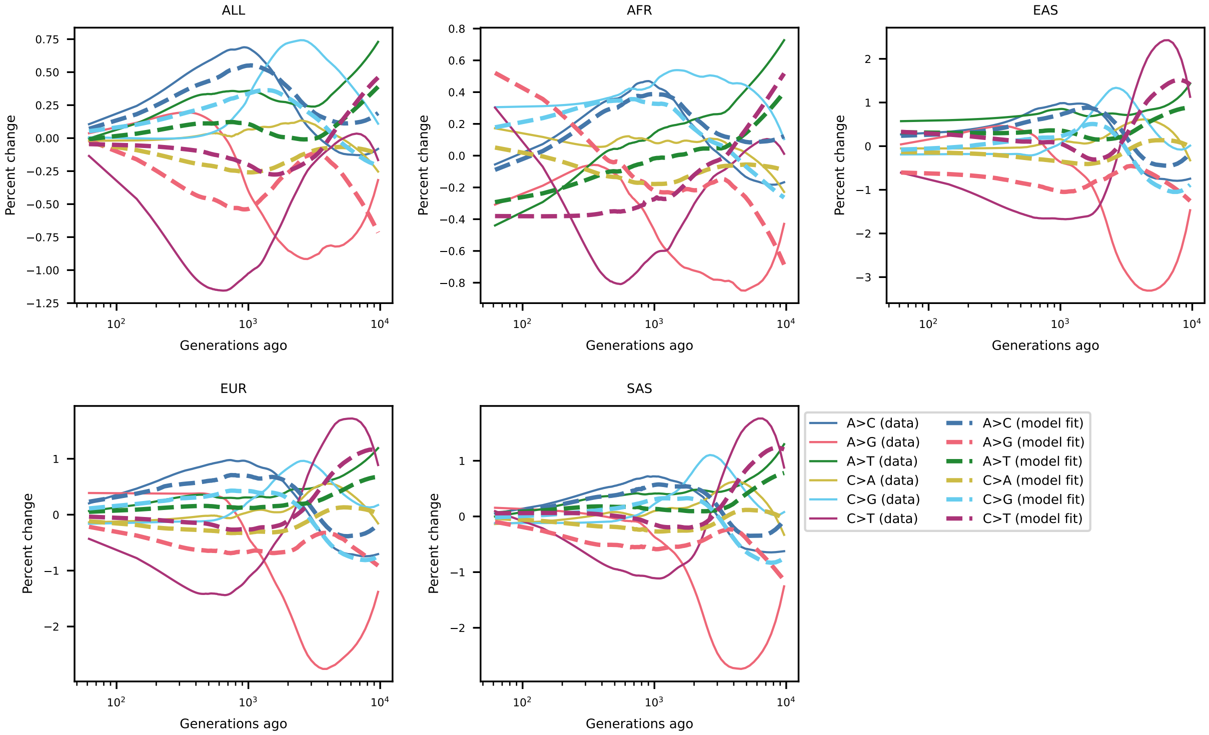


Figure S13: **Prediction of mutation spectrum history from** `tsdate`**-inferred generation times.**
